# KSHV upregulates the long non-coding RNA PURPL via p73 to suppress p53

**DOI:** 10.64898/2026.10.02.755609

**Authors:** Samia Showgan, Diana Meary, Guy Journo, Vyacheslav Gurevich, Meir Shamay

## Abstract

Long non-coding RNAs (lncRNAs) are a class of regulatory RNAs with lengths exceeding 200 nucleotides. The lncRNA PURPL (p53 upregulated regulator of p53 levels) is a p53-regulated gene. PURPL is an oncogenic lncRNA that represses p53 function; it is upregulated in many tumors and promotes cancer cell proliferation. Kaposi’s sarcoma-associated herpesvirus (KSHV) is the causative agent of all forms of Kaposi’s sarcoma (KS) and is tightly associated with primary effusion lymphoma (PEL). We found upregulation of PURPL in both de novo and chronically KSHV-infected cells. We found that p73 (a p53 homolog) associates with the PURPL promoter and upregulates PURPL expression. Previously, it has been shown that KSHV-encoded latency-associated nuclear antigen (LANA) stabilizes p73. We found that ectopic expression of LANA increased both p73 protein levels and PURPL expression. p73 knockdown abrogates PURPL upregulation following KSHV infection. Antisense oligonucleotides (ASO) targeting PURPL reduced PEL cell proliferation and increased acetylated p53 levels. In contrast to p53, p73 is not repressed by PURPL RNA. In summary, our study reveals how an oncogenic virus can utilize p73 to upregulate the oncogenic lncRNA PURPL as an additional way to repress p53. Moreover, we propose another mechanism by which p73 may promote tumor growth.

## Introduction

Long non-coding RNAs (lncRNAs) are a class of non-coding regulatory RNAs with lengths exceeding 200 nucleotides and are increasingly recognized for their critical roles in regulating disease processes, including cancer. The mammalian genome is transcribed into thousands of yet uncharacterized lncRNAs. Only a few lncRNAs have known biological functions, and in most cases, they are involved in the regulation of gene expression and function^1^. lncRNAs can contribute to the pathogenesis of different malignancies by promoting cell proliferation, migration, and invasion ^2^.

Viral infections are recognized as a causative factor in approximately 12% of all human cancer cases ^3,4^. Kaposi’s sarcoma-associated herpesvirus (KSHV) and Epstein-Barr virus (EBV) are two oncogenic viruses of the Herpesviridae family. KSHV is the causal agent of all forms of Kaposi’s sarcoma (KS) ^5^, and it is also tightly linked to other lymphoproliferative disorders, primary effusion lymphoma (PEL) ^6^ and Multicentric Castleman’s disease (MCD) ^7^, as well as a distinct cytokine-driven condition known as Kaposi sarcoma inflammatory cytokine syndrome (KICS)^8^. Similar to other herpesviruses, KSHV presents latent/dormant and lytic/productive phases. Most KSHV-associated malignant cells are latently infected, although a minority of lytic cells might contribute to the malignancy. Several lncRNAs, including MALAT1, H19, MEG3, NEAT1, UCA1, and KIKAT/LINC01061, have been shown to be dysregulated in KSHV-infected cells and tumors^9,10, 11^. In addition, ORF57 binds and regulates the expression of a subset of lncRNAs, including LINC00324, LINC00355 and LINC00839, which are involved in cell growth^12^. Another study found that KSHV-encoded miRNAs target 54 lncRNAs, including HOTTIP, DLEU2, HOTAIRM1, ANRIL, MEG3, and UCA1^13^.

The tumor suppressor p53 is a central regulator of cellular stress responses, playing essential roles in maintaining genomic stability by inducing cell cycle arrest, apoptosis, DNA repair, and senescence. Under normal conditions, p53 is kept at low levels through MDM2-mediated degradation^14^, but it is rapidly stabilized and activated in response to stress. Importantly, p53 function is frequently disrupted in cancer, either through mutation or viral interference. In the case of KSHV, several viral proteins have been shown to bind and repress p53 function, including the latency-associated nuclear antigen (LANA), viral interferon regulatory factor 3 (vIRF3/LANA2), vIRF1, and ORF45 ^15–19^.

p53 belongs to a family of transcription factors that include p63 and p73, which share structural similarities and overlapping target genes, but also have distinct roles in development and disease. While p53 is a well-established tumor suppressor, its family member p73 has a more complex and context-dependent role in cancer ^20^. Although p53 and p73 share structural similarities and in some cases overlapping transcriptional targets, their roles in tumorigenesis can differ. Unlike p53, which is frequently mutated across many cancer types, p73 mutations are detected in less than 0.5% of tumors^21,22^. The p73 family consists of two major subgroups, the transcriptionally active TAp73 isoforms and the ΔNp73 isoforms, which lack the N-terminal transactivation domain. Interestingly, overexpression of certain p73 isoforms has been found to have oncogenic properties in specific cancer contexts ^23–25^. The N-terminal-truncated ΔNp73 isoform shows oncogenic characteristics and is known for its dominant-negative repression of the TAp73 and p53 ^26^. Surprisingly, even the transcriptionally active TAp73 isoform, which is generally considered to have tumor-suppressive functions, can display oncogenic activity under certain conditions. In normal hepatocytes, only ΔNp73 is expressed, but most hepatocellular carcinoma cells express the full-length TAp73 ^27^. TAp73 has been shown to enhance prosurvival pathways, via the pentose phosphate pathway (PPP) or the activation of AP-1 ^28,29^. Moreover, it has been shown that TAp73 binds to p53 and inhibits p53-mediated transactivation following mild DNA damage, resulting in limited activation of p53 target genes to allow growth inhibition^30^.

Several studies that searched for lncRNAs upregulated during p53 activation detected a pronounced upregulation of the lncRNA LINC01021/RP11-46C20.1/LOC643401 ^31,32^. This lncRNA, later termed PURPL (p53 upregulated regulator of p53 levels), is upregulated in various colorectal cancer cells. PURPL reduces p53 levels by preventing the binding of p53 to its stabilizing protein Myb-binding protein 1A (MYBBP1A) ^33^. Subsequent research revealed its ability to promote various types of cancers, including melanoma^34^, liver^35^, ovarian^36^, breast^37^ and gastric^38^ cancers. It has been shown that silencing PURPL significantly reduced melanoma cell viability, colony formation, migration, and invasiveness^39^. Moreover, PURPL is upregulated during senescence^40^, and enhances the radioresistance of cancer cells and therefore might serve as a therapeutic target for improved cancer treatment ^41^.

In this study, we identified PURPL as an upregulated lncRNA following KSHV infection of epithelial, endothelial, and B cells, as well as KSHV-associated PEL cells. We found that the p53 homolog p73 directly upregulates PURPL expression. p73 is necessary for KSHV-induced PURPL upregulation, as PURPL upregulation is abolished upon p73 knockdown. A previous study has shown that KSHV latent protein LANA stabilizes p73, leading to its accumulation in KSHV-infected cells ^42^. In agreement with this study, we found that LANA expression leads to p73 accumulation and PURPL upregulation. In addition, we found that PURPL reduces p53 acetylation. Interestingly, while PURPL reduces p53 levels, it does not negatively regulate p73 levels. Therefore, we propose that p73 can negatively regulate p53 levels via PURPL upregulation. Knockdown of PURPL in KSHV-associated primary effusion lymphoma cells inhibits cell growth. In summary, we revealed a novel mechanism by which p73 inhibits the tumor suppressor p53 through the lncRNA PURPL. This mechanism may explain the overexpression of p73 in certain tumors. Oncogenic viruses, such as KSHV and EBV, exploit PURPL as an additional mechanism to inhibit p53 and promote cell growth.

## Results

### PURPL is upregulated in KSHV-infected cells

In a previous study^43^, we performed RNA-sequencing on two KSHV-associated primary effusion lymphoma (PEL) cell lines, BC3 and BCBL1, and a KSHV-negative lymphoma cell line BJAB, a well-established control in the field. To determine the effect of KSHV on cellular gene expression following infection, a comparison between BJAB and BJAB infected with a recombinant KSHV virus (BJAB.219) was performed. Reanalysis of these data revealed a dramatic upregulation of the lncRNA PURPL (also known as LINC01021, LOC643401, and RP11-46C20.1) in both PEL cell lines and BJAB.219 (Fig. 1A) compared to uninfected cells. We validated PURPL upregulation in the B-cell lines BC3 and BCBL1 (Fig. 1B) and BJAB.219 (Fig. 1C) via RT-qPCR. We also observed upregulation of PURPL in endothelial (TIME) and epithelial (SLK, HEK293, and HCT116) cells infected with KSHV (Fig. 1D-G). Interestingly, PURPL was also upregulated in cells infected with the related gammaherpesvirus, Epstein-Barr virus (EBV) (Fig. S1A). In summary, we found upregulation of PURPL in cells infected with KSHV and EBV.

**Figure 1.**
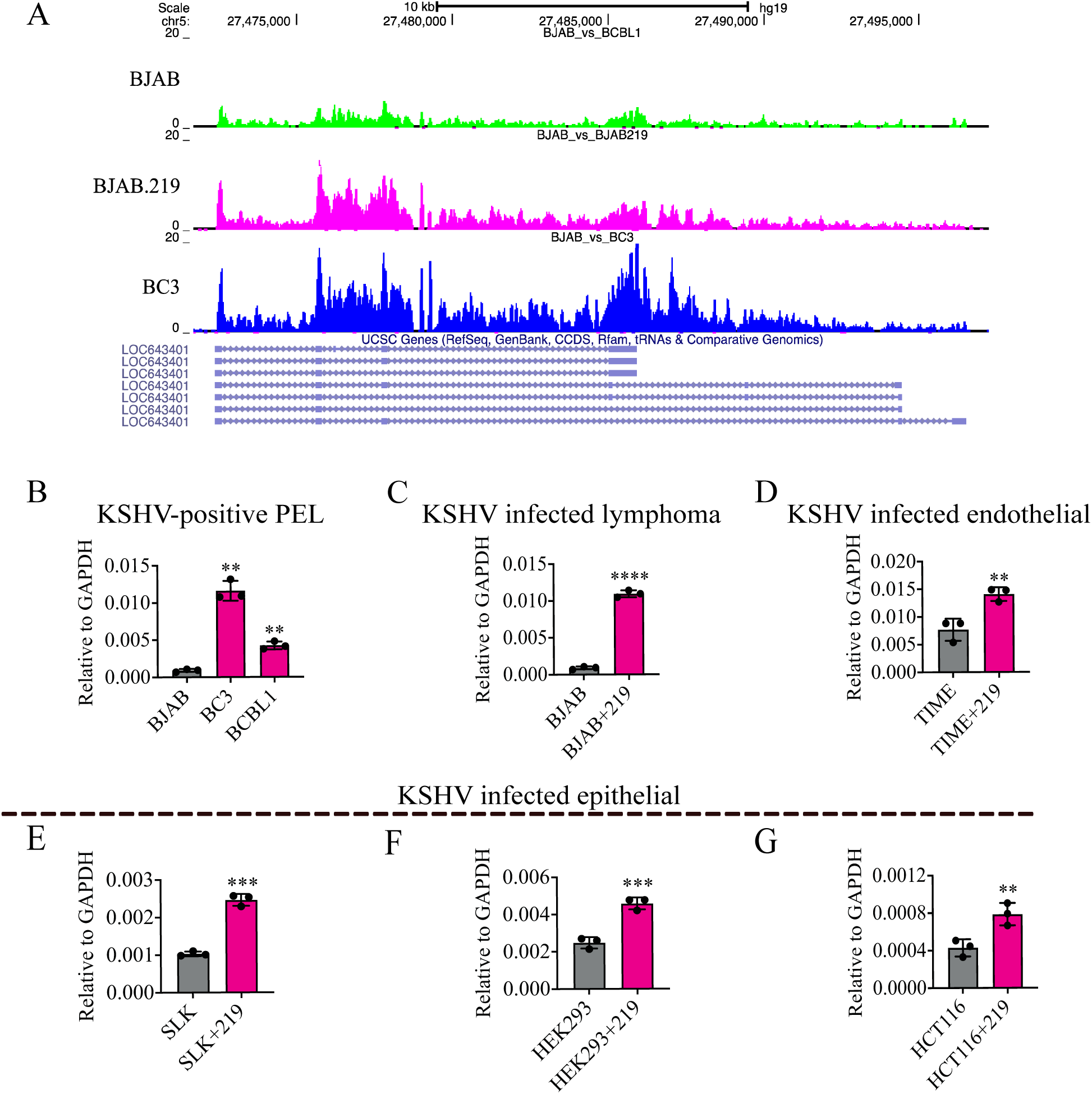
The lncRNA PURPL is upregulated in KSHV-infected cells. (A) Reanalysis of RNA-seq from Journo et al ^43^, for the expression of PURPL (LOC643401) in KSHV-infected cells BC3 and de novo infected BJAB.219 compared to the non-infected lymphoma BJAB and visualized on the UCSC genome browser ^61^. (B) RT-qPCR analysis for PURPL expression on RNA isolated from PEL (BC3, BCBL1), (C) de novo KSHV-infected B-cells (BJAB.219), (D) endothelial (TIME) and (E) epithelial SLK, (F) HEK293 and (G) HCT116. All results were normalized to GAPDH. Experiments were performed in three biological replicates (n = 3), and data are represented as means ± SD. One-tailed T-tests were performed. **=p<0.01 ***=p<0.001 ****=p<0.0001.

### TAp73 isoforms activate PURPL promoter

PURPL was identified as one of the most upregulated lncRNAs upon DNA damage and p53 activation following Nutlin-3 treatment^31,32^. p53 directly increases PURPL transcription via binding to a p53 consensus binding site within the PURPL promoter. Surprisingly, we observed PURPL upregulation in BJAB cells expressing a mutant (non-functional) p53 protein ^44^. The fact that the virus appears to exploit a p53-independent pathway to regulate PURPL expression prompted us to investigate how KSHV regulates PURPL.

To study the regulation of PURPL by KSHV, we cloned the PURPL promoter upstream of a luciferase reporter gene (Fig. 2A). Mutation of the p53 consensus binding site within PURPL promoter significantly reduced PURPL promoter activity in KSHV-infected BC3 cells (Fig. 2B), and BJAB.219 (expressing mutant p53) (Fig. 2C), suggesting that this binding site is required for PURPL upregulation independent of a functional p53.

**Figure 2.**
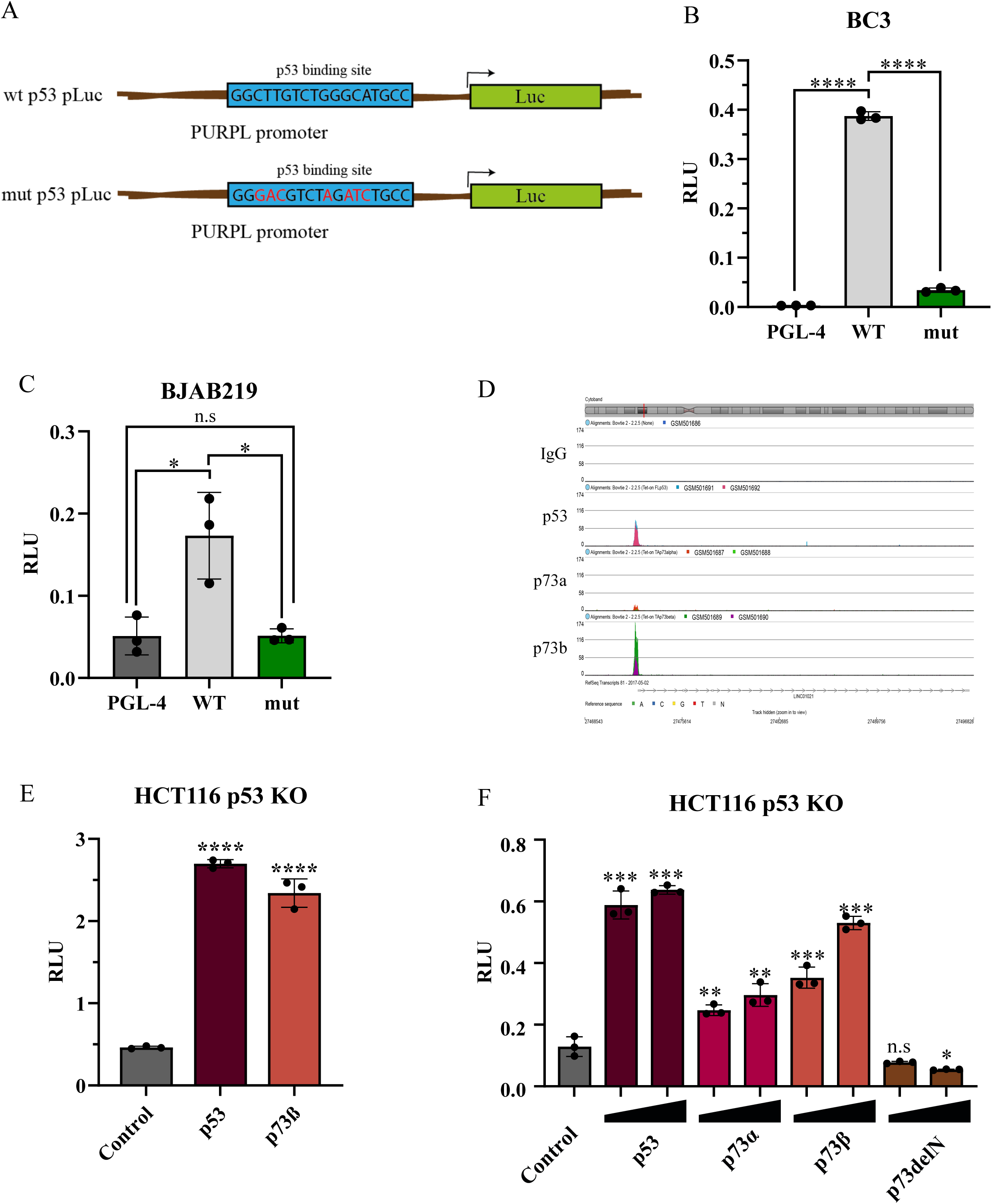
TAp73 isoforms activate PURPL promoter. (A) Schematic representation of WT and mutant PURPL promoter luciferase reporter plasmids. (B-C) BC3 and BJAB219 cells were transiently transfected with the empty Luc vector (control), wt, or mutant luc-PURPL promoter construct together with control (Renilla) reporter, and dual luciferase reporter assay was performed. (D) Re-analysis of published ChIP-seq datasets for p53, TAp73α, and TAp73β (GEO: GSE15780) generated in p53-null Saos-2 cells stably expressing each protein individually, to assess their binding at the PURPL promoter. (E) HCT116 p53 KO cells were transfected with luc-PURPL promoter together with expression vectors for p53 and p73β (15 ng each), and dual luciferase reporter assay was performed. (F) same as in E, but expression vectors for p53 and different p73 isoforms (5 ng and 15 ng) were transfected. Luciferase values are shown as relative luciferase units (RLU) normalized to Renilla luciferase internal control. Experiments were performed in three independent biological replicates (n = 3), and data are represented as means ± SD. Values in the graph indicate P values from 2-tailed T-tests. (ns) denotes non-significant values.

The DNA-binding domains (DBDs) of the two p53 homologs, p63 and p73, are 60% and 63% identical to p53, respectively. Consistent with this, many p53-responsive promoters are transactivated by p73α and p73β ^45,46^. The p73 family comprises two major subgroups: the transcriptionally active TAp73 isoforms, p73α and p73β, and the ΔNp73 isoforms, which lack the N-terminal transactivation domain (Fig. S2A). To determine whether p73 binds to the PURPL promoter, we re-analyzed publicly available ChIP-seq datasets generated in p53-null Saos-2 cells engineered to stably express p53, TAp73α, or TAp73β individually (GEO: GSE15780)^47^ (Fig. 2D). In addition, we reanalyzed p63 ChIP-seq data generated in ME-180 cells, overexpressing p63 (GEO: GSE135257)^48^ (Fig. S2B). We found a strong association of both p53 and p73β with the PURPL promoter.

Next, we co-transfected p53, p73 or p63 expression plasmids together with the PURPL luciferase reporter into HCT116 p53 knock-out (p53 KO) cells to ensure that the observed transactivation was independent of endogenous p53 and performed a luciferase assay. Since only p73β was strongly enriched at the PURPL promoter in the ChIP-seq, we therefore first examined it. We found that p73β activated PURPL promoter to a similar level as p53 (Fig. 2E). We then examined additional p73 isoforms (Fig. 2F) and found that both p73α and p73β upregulated PURPL promoter activity, whereas ΔNp73 repressed it. In contrast, p63α showed only a mild activation (1.32-fold) of the PURPL promoter, while ΔNp63 slightly repressed it (Fig. S2C). Together, these findings support a role for TAp73 isoforms in the transcriptional activation of PURPL.

### The upregulation of PURPL by p73 is dependent on the p53 consensus binding site within PURPL promoter

To assess whether p73 can regulate endogenous PURPL, we used H1299 cells (p53-deficient) with a doxycycline-inducible p73α (TET-on) system^49^. We confirmed the induction of p73α expression following doxycycline treatment (Figure 3A) and found that PURPL was significantly upregulated following p73 expression (Figure 3B). To test whether p73 activates the PURPL promoter via the p53 consensus binding site we performed luciferase reporter assay with the mutated PURPL promoter in HCT p53 KO cells. This experiment revealed that the mutation at the p53-binding site abrogated the ability of p73α and p73β to induce PURPL expression (Fig. 3C). These experiments confirmed that p73 upregulates PURPL levels directly via the p53 consensus binding site within the PURPL promoter.

**Figure 3.**
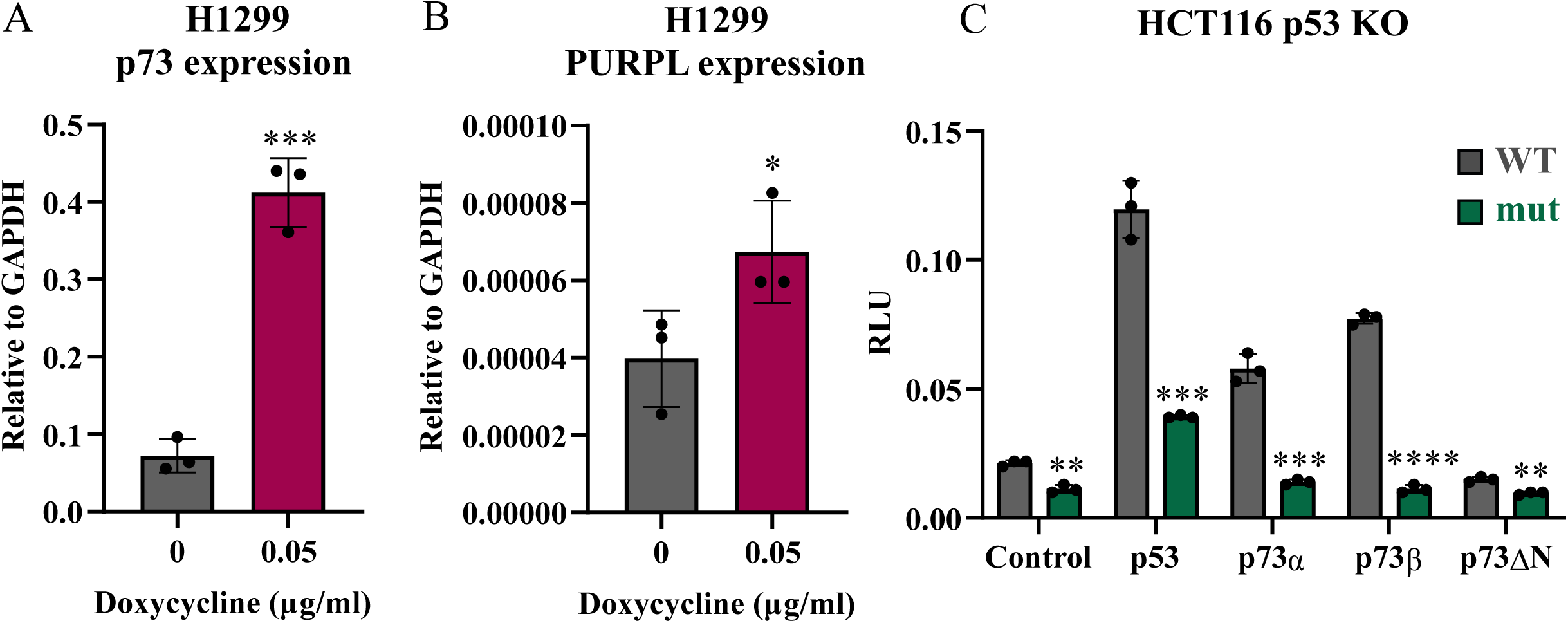
The upregulation of PURPL by p73 is dependent on the p53 consensus binding site within PURPL promoter. (A-B) H1299 cells with p73-TET-ON system kept untreated or treated with doxycycline for 24 h, and RNA was isolated and subjected to RT-qPCR analysis for p73α (A) and PURPL (B) expression. (C) HCT116 p53 KO cells were transfected with wild-type (WT) or mutant (Mut) PURPL promoter constructs in the presence of p53, p73α, p73β, or ΔNp73α expression vectors. Experiments were performed in three biological replicates (n = 3) and data are represented as means ± SD. Luciferase data are presented as relative luciferase units (RLU) normalized to Renilla luciferase internal control. RT-qPCR results are normalized to GAPDH expression.

### KSHV LANA promotes PURPL expression

A previous study has shown that KSHV-encoded LANA can interact with p73α and ΔNp73, stabilizing their protein levels ^42^. Since we found that p73 upregulates PURPL expression, we next investigated whether LANA contributes to PURPL upregulation. To this end, we generated BJAB cells expressing doxycycline-inducible LANA. Upon doxycycline treatment, LANA expression was induced, as confirmed by western blot and RT-qPCR analysis (Fig. 4A-B). LANA expression led to increased p73 protein levels (Fig. 4A) and PURPL RNA expression compared to control cells (Fig. 4C), suggesting that LANA is sufficient to increase p73 and PURPL levels in BJAB cells.

**Figure 4.**
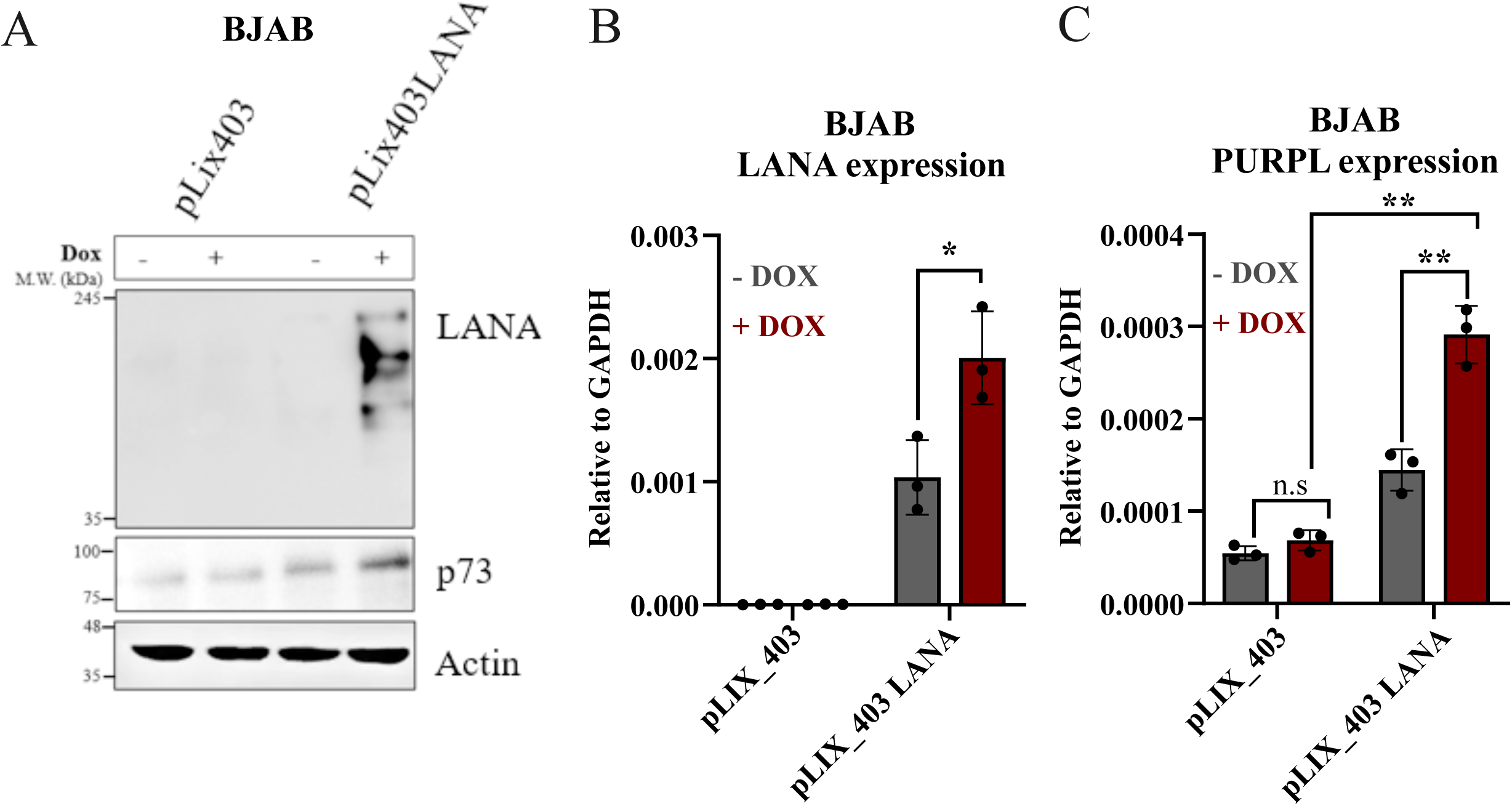
KSHV LANA promotes PURPL expression. (A) Western blot analysis of p73 and LANA protein levels in BJAB cells transduced with doxycycline-inducible pLIX403_LANA or empty pLIX403 vector following doxycycline treatment. Actin served as a loading control. (B-C) RT-qPCR analysis of LANA (B) and PURPL (C) in the same cells. Relative expression levels were normalized to GAPDH. RT-qPCR experiments were performed in three biological replicates (n = 3) and data are represented as means ± SD.

### p73 knockdown reduces PURPL expression in KSHV-infected and uninfected cells

In agreement with Santag et al^42^, we found increased p73 protein levels in PEL cell lines (BC3 and BCBL1) and infected BJAB.219 compared to BJAB (Fig. 5A). In contrast to p73, the expression of p63 protein was not consistent with KSHV infection, as we were able to detect it only in BC3 cells (Fig. S3A). To test the role of p73 during PURPL upregulation following KSHV infection, we knocked down (KD) p73 in BJAB cells. The cells were transduced with lentivirus expressing a control shRNA or different shRNAs targeting p73. Three shRNAs targeting p73 were confirmed to efficiently knockdown p73 by Western blotting (Fig. 5B). Based on sequence analysis of the shRNA target sites, two of the shRNAs target regions common to both TAp73 and ΔNp73 isoforms (shRNA 3 and 4), whereas one shRNA targets a region present only in the TAp73 transcripts and should not affect ΔNp73 (shRNA 1). The cells were then infected with recombinant KSHV BAC16, and successful infection was validated by detecting LANA RNA and protein via RT-qPCR and Western blotting, respectively (Fig. 5C-D). Consistent with our previous observations, KSHV infection upregulated PURPL expression. Interestingly, this PURPL upregulation was abolished in p73 KD cells for all three p73 shRNAs (Fig. 5E).

**Figure 5.**
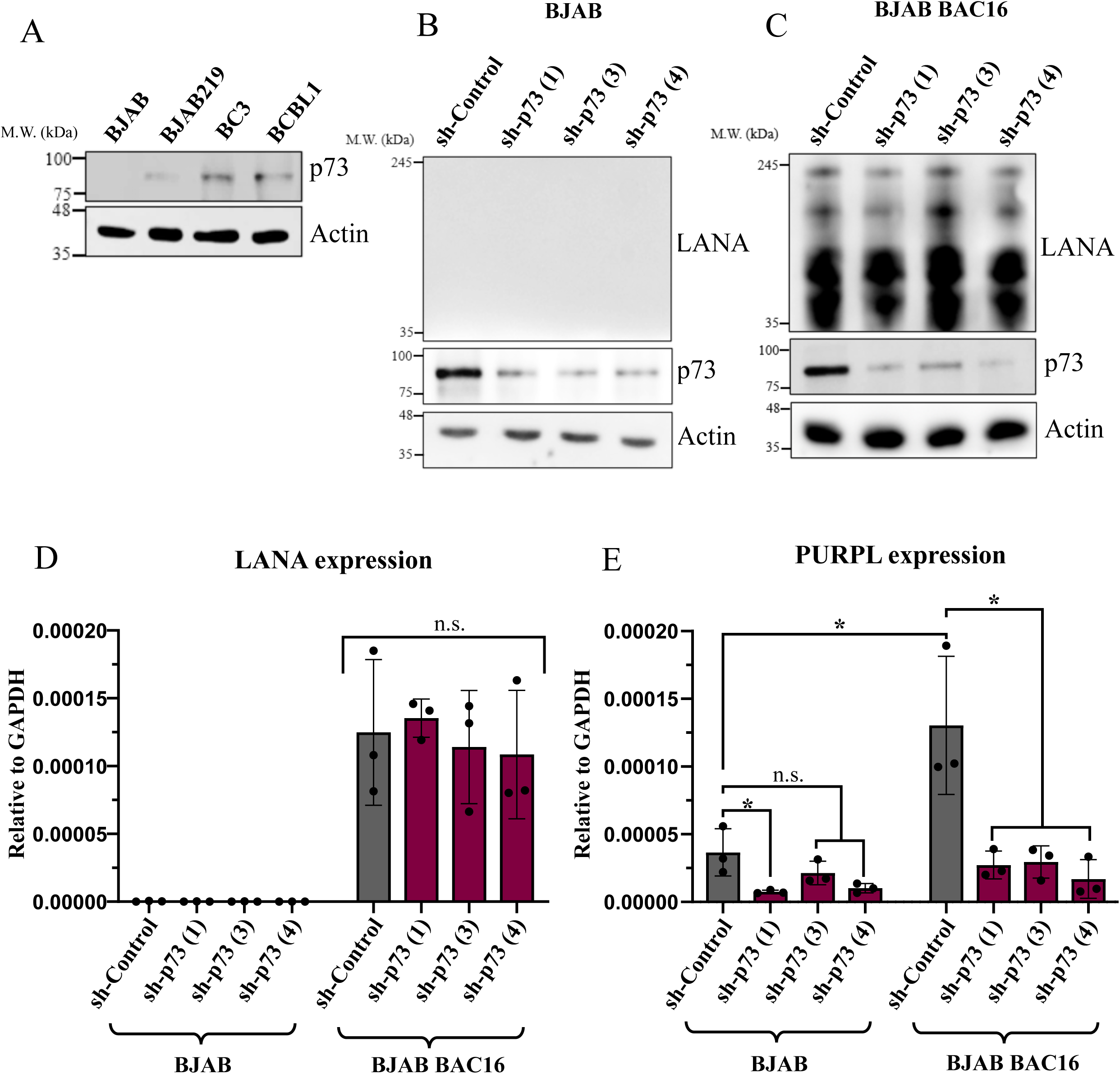
p73 knockdown abolishes PURPL upregulation in KSHV-infected cells. (A) Western blot analysis of TAp73 protein levels in uninfected BJAB and KSHV-infected cells (BJAB.219, BC3, and BCBL1). (B–C) BJAB (B) and BJAB-BAC16 (C) cells were transduced with control shRNA (sh-Control) or three different shRNAs targeting p73. Western blot analysis of TAp73 and LANA protein levels is shown. Actin was used as a loading control for the Western blot experiments. (D–E) RT-qPCR analysis of LANA (D) and PURPL (E) RNA expression in the same cells. RT-qPCR experiments were performed in three biological replicates (n = 3) and data are represented as means ± SD.

Since a previous study reported that p73 knockdown in U2OS osteosarcoma cells reduces cell proliferation^50^, we repeated the p73 shRNA knockdown in U2OS cells, which harbor wild-type p53, to examine whether PURPL expression is also affected in this system. The knockdown of p73 was validated by Western blot analysis (Fig. 6A) and qPCR (Fig. 6B). Depletion of p73 led to significantly lower PURPL levels (Fig. 6C), indicating that p73 also upregulates PURPL in uninfected cells. Our data indicate that although PURPL is a known p53-responsive lncRNA, the reduction of PURPL expression following p73 knockdown in U2OS cells suggests that p73 contributes to PURPL regulation even in the presence of wild-type p53.

**Figure 6.**
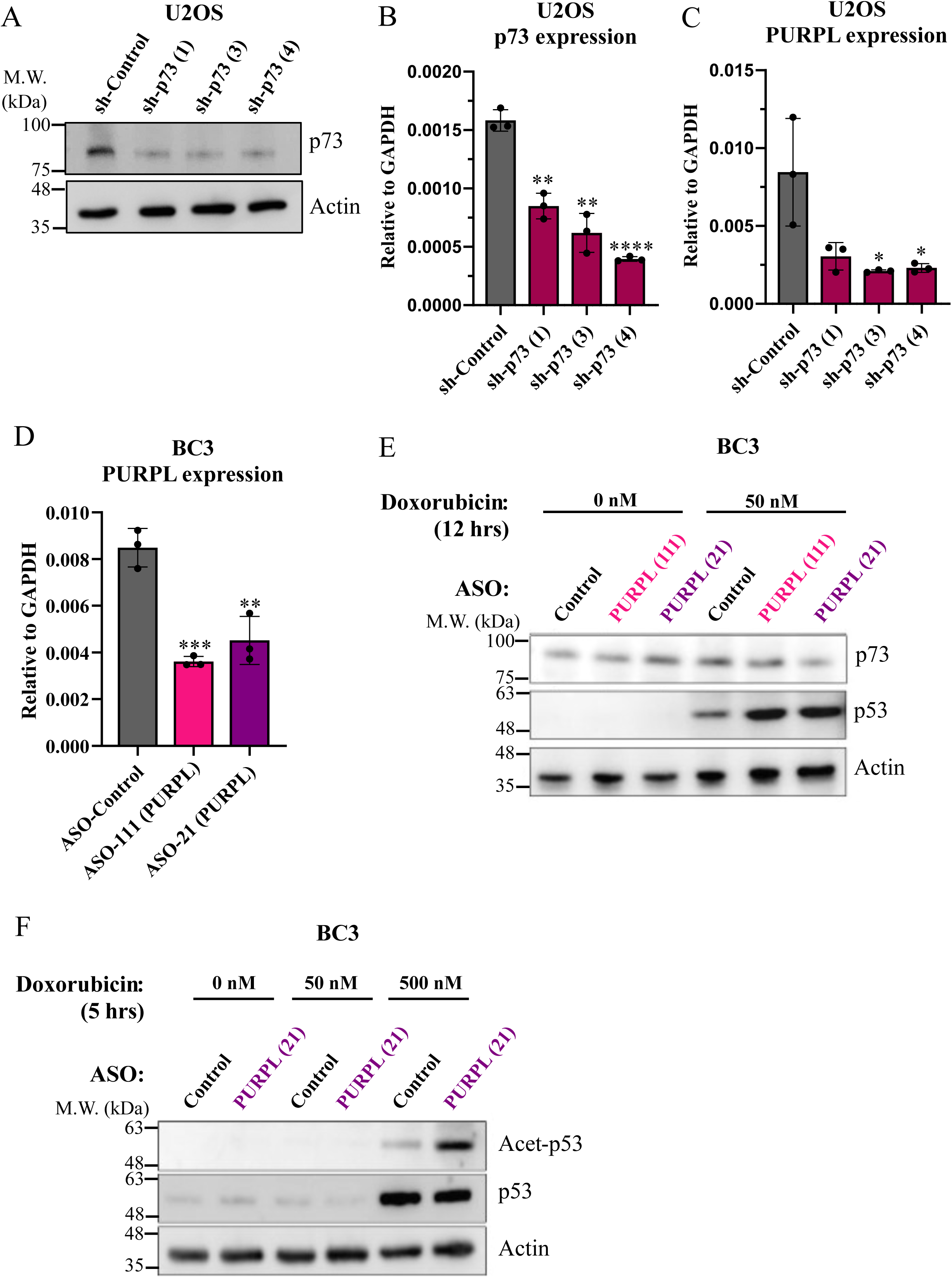
PURPL is a negative regulator of p53 levels and acetylation, but not p73. (A) Western blot analysis of TAp73 protein levels in U2OS cells following transduction with the indicated shRNAs targeting p73. (B–C) RT-qPCR analysis of p73 (B) and PURPL (C) RNA levels in U2OS cells following p73 knockdown. (D) RT-qPCR analysis of PURPL expression in BC3 cells treated with control antisense oligonucleotide (ASO-Control) or PURPL-targeting ASOs (ASO-21 and ASO-111). RT-qPCR experiments were normalized to GAPDH, were performed in three biological replicates (n = 3) and data are represented as means ± SD. (E) Western blot analysis of TAp73 and p53 protein levels in BC3 cells treated with control ASO or PURPL-targeting ASO (ASO-21 and ASO-111) for 72 hours, followed by 12 hours of doxorubicin treatment (0 or 50 nM). Actin was used as a loading control. (F) Western blot analysis of TAp73 and acetylated p53 protein levels in BC3 cells treated with control ASO or PURPL-targeting ASO (ASO-21) for 48 hours, followed by 5 hours of doxorubicin treatment (0, 50 or 500 nM). Actin was used as a loading control for the Western blot experiments.

### PURPL is a negative regulator of p53 levels and acetylation, but not p73

It has been shown that PURPL negatively regulates p53 protein levels^33^. Therefore, we investigated whether p73 is also regulated by PURPL levels. We designed two antisense oligonucleotides (ASOs) targeting PURPL, ASO-111 and ASO-21. PURPL knockdown was performed using ASOs, as antisense-mediated RNase H degradation is more efficient for targeting nuclear long noncoding RNAs such as PURPL than RNAi-based approaches like shRNA. We knocked down PURPL using these ASOs in the BC3 cell line, which expresses WT p53 and high levels of PURPL (Fig. 1B). Both ASO-111 and ASO-21 showed PURPL knockdown (Fig. 6D). We then added the DNA-damage inducer doxorubicin for 12 hours and determined the levels of p53 and p73 by Western blot analysis (Fig. 6E). While p53 levels increased as expected following PURPL KD, p73 did not accumulate, suggesting that p73 is not sensitive to PURPL levels.

A previous study found that PURPL interferes with p53 binding to MYBBP1A, leading to reduced p53 levels^33^. MYBBP1A is important for the interaction between p53 and the co-activator CBP/p300^51^. Since CBP/p300 mediates p53 acetylation, a necessary step in p53 activation, we hypothesized that PURPL inhibits p53 acetylation. We found that long-term (12 h) doxorubicin treatment leads to increased p53 levels in PURPL KD cells (Fig. 6E), whereas short-term (5 h) doxorubicin treatment enhances p53 acetylation in PURPL KD cells (Fig. 6F). This suggests that following short-term DNA damage, PURPL decreases the acetylation levels of p53.

### PURPL knockdown results in reduced proliferation of PEL cells without increasing apoptosis

Several studies suggest that PURPL is an oncogenic lncRNA that promotes cancer cell proliferation, and its knockdown results in cell growth inhibition^33,52,53^. To test if PURPL expression is essential for the proliferation of KSHV-positive cells, we monitored cell proliferation using an automated cell counter following PURPL KD. Our results demonstrate a profound inhibition of cell growth in BC3 cells following PURPL KD (Fig. 7A). To further assess the percentage of viable cells based on metabolic activity, we analyzed cells from the same ASO-transfected populations by XTT assay at the final time point (120 h post-transfection). Both ASO-21 and ASO-111 significantly reduced the percentage of viable cells compared with the control ASO (Fig. 7B), consistent with the reduced cell growth observed in the proliferation assay (Fig. 7A). To determine whether this inhibition of cell growth is due to increased apoptosis after PURPL knockdown, flow cytometry was performed after staining cells with propidium iodide (PI) to detect cell death and Annexin V to detect early apoptosis. We found that in KSHV-infected PEL cells, reduced cell proliferation after PURPL KD is not associated with higher levels of cell apoptosis or cell death (Fig. 7C, S4A-B). These results indicate that the reduced XTT signal likely reflects the lower number of metabolically active cells resulting from impaired proliferation, rather than increased cell death. Together, these findings indicate that KSHV induces p73 expression, thereby upregulating PURPL, which in turn inhibits p53 activity and may contribute to viral oncogenesis.

**Figure 7.**
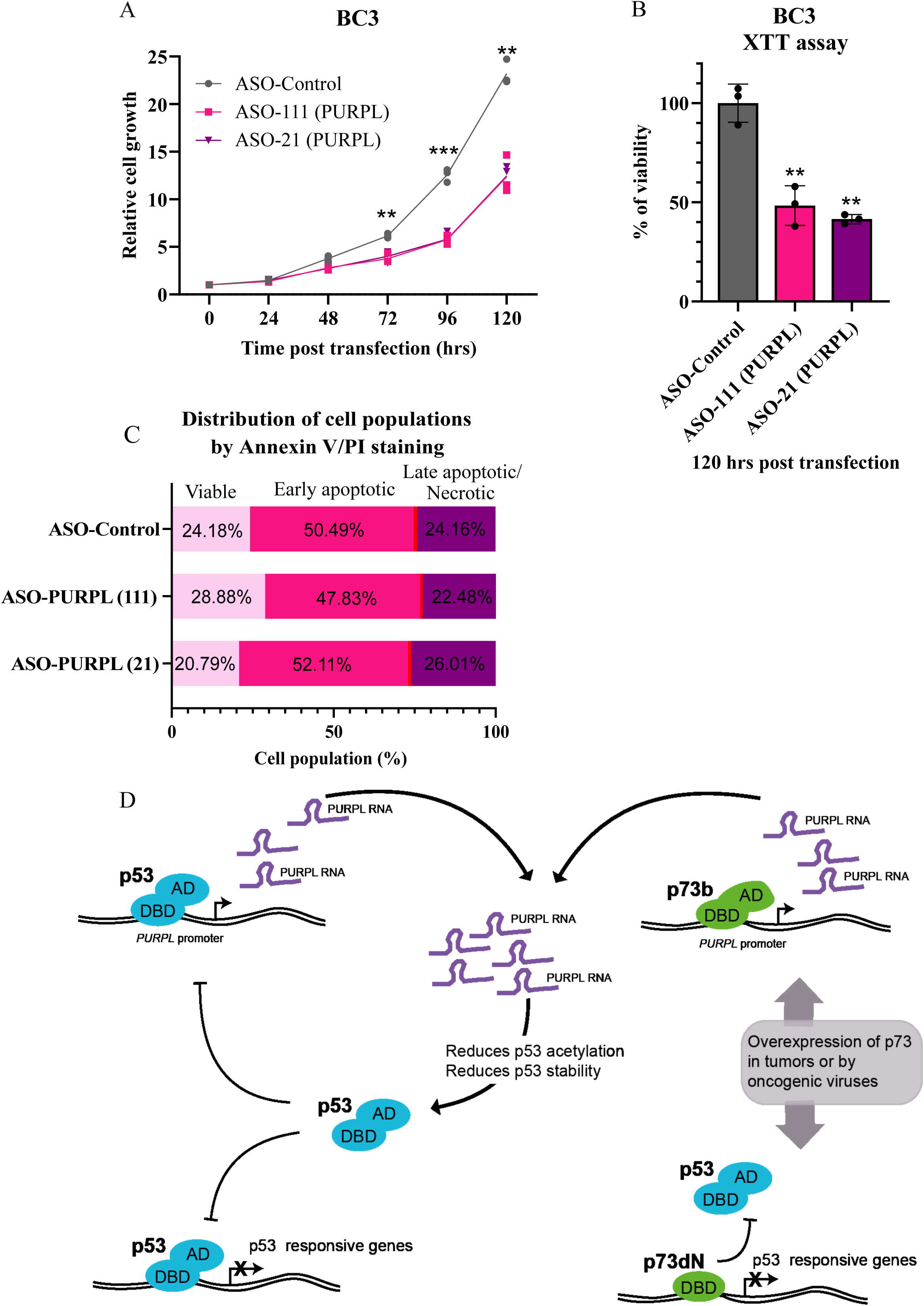
PURPL knockdown reduces PEL cell proliferation without increasing apoptosis. (A) BC3 cells transfected with ASO-Control or ASO-PURPL (ASO-21 and ASO-111) were counted every 24 h via an automated cell counter to evaluate proliferation. The cell proliferation assay was performed in three biological replicates (n = 3), and data are represented as means ± SD. (B) A population of the BC3 cells used in the proliferation assay shown in Figure A were subjected to an XTT assay at 120 hours post-transfection. Percentage of cell viability was normalized to ASO-Control. The XTT assay was performed in three biological replicates (n = 3), and data are represented as means ± SD. (C) Flow cytometry analysis of apoptosis and cell death in BC3 cells treated with ASO-Control or ASO-PURPL (ASO-21 and ASO-111). Cells were stained with Annexin V to detect early apoptosis and propidium iodide (PI) to detect late apoptosis and cell death. (D) Schematic illustration of the proposed model regarding the inhibition of p53 by p73.

## Discussion

PURPL has been identified as a p53-regulated lncRNA; however, oncogenic viruses, such as KSHV, can inhibit p53 activity. In addition, we found PURPL upregulation following KSHV infection in BJAB cells expressing a mutant p53 protein. Therefore, an alternative, p53-independent pathway likely controls PURPL expression in this case. A previous study^54^ reported PURPL upregulation in another p53-mutant cell line, DLD1, which supports our observation. To our surprise, a mutation in the p53 consensus binding site within the PURPL promoter abrogated PURPL upregulation in KSHV-infected cells, suggesting that the factor regulating PURPL in these cells binds the p53 consensus site. Reanalysis of available ChIP-seq data revealed a strong association of p73β with the PURPL promoter. p73β can upregulate PURPL, and a mutation in the p53 consensus binding site abrogates this activity. Another lncRNA shown to be regulated by p73 is FER1L4, but unlike PURPL, p73-mediated regulation of FER1L4 promotes cell cycle arrest and apoptosis^55^. p73 is upregulated (similar to PURPL) in many tumors, suggesting that p73 may promote oncogenic functions in certain tumor contexts. The N-terminal deleted isoform (ΔNp73) can serve as a dominant negative for p53 by binding DNA at p53 consensus binding sites and thereby competing with p53 binding, and, due to the lack of an activation domain, prevents transcription activation. Accordingly, ΔNp73 exhibits oncogenic characteristics, as it can inhibit the pro-apoptotic activity of p53 and block its function as a tumor suppressor ^56^. Therefore, the current model is that p73 can repress p53 activity via its N-terminally truncated form. A negative regulator of p53 that is upregulated by p73 is MDM2, but in this case, MDM2 binds and represses p73’s ability to activate transcription ^56^. Our study provides an additional mechanism by which p73 can repress p53 activity: by upregulating PURPL, leading to reduced p53 acetylation and protein levels (Fig. 7D). Since PURPL is upregulated following KSHV infection, this pathway may contribute to suppression of p53 activity in infected cells through reduced p53 acetylation. In this respect, it is important to note that, unlike p53, we found that p73 is not negatively regulated by PURPL. What is the connection between p73, KSHV, and PURPL? A previous study^42^ has shown p73 upregulation in KSHV-infected cells. They have found that the KSHV-encoded LANA protein interacts with p73, leading to its accumulation. Our findings further support this model, as ectopic LANA expression was accompanied by elevated p73 protein levels and increased PURPL expression. We found that PURPL upregulation in KSHV-infected cells is mediated by p73, since PURPL upregulation following KSHV infection was abolished in p73 KD cells. PURPL upregulation contributes to PEL cell growth, as PURPL KD leads to inhibition of PEL cell proliferation. In summary, our study shows that p73 plays a key role in KSHV pathogenesis by upregulating PURPL levels, thereby inhibiting p53 and promoting tumor growth. This highlights PURPL as a promising marker and therapeutic target in KSHV-associated malignancies.

## Materials and Methods

### Antibodies

Primary antibodies included anti-p73 rabbit polyclonal antibody (ab14430; Abcam), anti-p53 rabbit monoclonal antibody (#2527; Cell Signaling Technology), anti-acetyl-p53 rabbit polyclonal antibody (#2525S; Cell Signaling Technology), anti-p63 rabbit monoclonal antibody (ab124762; Abcam), anti-β-actin mouse monoclonal antibody (A2228; Sigma-Aldrich), and anti-LANA rat monoclonal antibody (MABE1109; Sigma-Aldrich). Secondary antibodies included donkey anti-rabbit HRP-conjugated antibody (NA934V; Amersham), goat anti-rat HRP-conjugated antibody (112-035-062; Jackson ImmunoResearch), and sheep anti-mouse HRP-conjugated antibody (NA931V; Amersham).

### Cell lines

BC3, BCBL1 and BJAB cells (kindly provided by Richard F. Ambinder), BJAB pLIX403, BJAB pLIX403_LANA, BJAB.219, and BJAB.BAC16 were cultured in RPMI-1640 medium supplemented with 20% FBS (fetal bovine serum), 2 mM L-glutamine, penicillin-streptomycin (100 IU/mL and 100 μg/mL, respectively) and 1 mM Sodium-Pyruvate. K562, BC-1, RAJI, AKATA and p73α TET-on H1299 cells (kindly provided by Yosef Shaul) were maintained in RPMI-1640 medium supplemented with 10% FBS and the same supplements as the PEL cell lines. HCT116 p53-KO and HCT116 (kindly provided by Bert Vogelstein) and HCT116.219 were cultured in McCoy’s 5A medium supplemented with 10% FBS and the same supplements as the PEL cell lines. HEK293, HEK293.219, HEK293T, U2OS (kindly provided by Yosef Shaul), Vero-rKSHV.219^57^ (kindly provided by Jeffrey Vieira) and iSLK BAC16 mCherry^58^ (kindly provided by Ronit Sarid), a version of BAC16^59^, were maintained in DMEM medium supplemented with 10% FBS and the same supplements as the PEL cell lines. TIME and TIME.219 cells were maintained in Vascular Cell Basal Medium (VCBM) supplemented according to the manufacturer’s (ATCC) guidelines. Cell lines infected with rKSHV.219 were maintained under 1 µg/mL puromycin selection. iSLK cells infected with the BAC16 mCherry virus were maintained in 1200 µg/mL hygromycin, while BJAB cells infected with the same virus were maintained in 200 µg/mL hygromycin. Unless stated otherwise, the H1299 inducible cells were treated with 0.05 µg/mL doxycycline (Dox) for 24 hours to induce p73α expression. BJAB cells transduced with pLIX403 or pLIX403_LANA lentiviral vectors were similarly treated to induce LANA expression. All cell lines were grown in 5% CO_2_ at 37°C.

### Plasmids

To generate the PURPL-luc construct, primers for the PURPL promoter were designed and used for PCR amplification (PURPL promoter F and R; listed in Table S1). The PCR product from the PURPL amplification was cloned into pGEM-T Easy vector and subcloned into the pGL4.11[luc2P] (#<u>E6661</u>, Promega) vector using the EcoRV and NcoI restriction sites. A mutant PURPL promoter harboring substitutions in the p53 binding site was generated using the Q5 Site-Directed Mutagenesis Kit (New England Biolabs, E0554S) according to the manufacturer’s instructions. Mutagenic primers (Q5SDM F and R; listed in Table S1) were used to substitute seven nucleotides within the p53 binding site (mutated sequence listed in Table S2). Expression plasmids encoding p53, p73α, p73β, and p73αΔN(1–61) were kindly provided by Yosef Shaul. Plasmids encoding p63α and p63αΔN were kindly provided by Moshe Oren. pLIX_403 was a kind gift from David Root (Addgene, # 41395), and pLIX_LANA was previously described^60^.

### Antisense oligonucleotide (ASO) treatment

KSHV-infected PEL cells (BC3) were transfected using the Lipofectamine reverse transfection protocol with either ASO-Control, ASO-21, or ASO-111 at a final concentration of 50 nM. Transfections were performed for 48 or 72 hours, as indicated. For proliferation assays, apoptosis analysis, and Western blot experiments, BC3 cells were seeded at 3 × 10⁵ cells per well and reverse-transfected with ASO-Control, ASO-21, or ASO-111 (50 nM). For cell proliferation experiment, cell counts were measured every 24 hours for a total of 120 hours using an automated cell counter. In indicated experiments, cells were treated with doxorubicin (50 or 500 nM) for 5 or 12 hours prior to harvest. Oligonucleotide sequences are listed in Supplementary Table S1.

### Quantification of viable cells by XTT assay

To assess the percentage of viable cells, the XTT Cell Proliferation Kit II (#11465015001, Roche) was used according to the manufacturer’s instructions. Briefly, following transfection of BC3 cells with the indicated ASOs, 5,000 cells from each treatment group were seeded per well in a clear 96-well plate. At 120 hours post-transfection, the final time point of the cell-counting experiment, the XTT assay was performed. A mixture containing 49 µl XTT labeling reagent and 1 µl electron-coupling reagent was added to each well, followed by incubation for 4 h at 37°C in 5% CO₂. Absorbance was measured at 490 nm, and 630 nm was used as the reference wavelength. Blank wells containing complete growth medium and XTT reagent mixture without cells were included for background subtraction.

### Generation of de novo infected cells

The iSLK BAC16 mCherry cells were reactivated with 1 mM sodium butyrate (NaB) together with 1 μg/mL doxycycline for 24 hours. The medium was then removed, and BJAB cells were added for coculture with RPMI 1640 medium. After 3 days, infected BJAB cells were transferred to a new flask and treated with 200 μg/mL hygromycin for selection and maintenance. rKSHV.219^57^ virus was generated from Vero.219 cells. Vero.219 cells were induced with 1.25 mM NaB and 20 ng/mL phorbol ester (TPA) in DMEM for 24 hours. After induction, cells were washed twice with PBS and overlaid with DMEM containing 20 ng/mL TPA. Supernatants were collected every 2–3 days based on RFP expression. This cycle was repeated until 240 mL of supernatant was collected. Supernatants were centrifuged at 600 × g for 7 minutes at 4°C, filtered through 0.45 µm filters, and centrifuged overnight at 15,000 × g at 4°C to pellet the virus. Pellets were resuspended in 1 mL serum-free DMEM and incubated at 4°C overnight. The resuspended virus was aliquoted and stored at –80°C. Target cells were seeded in 6-well plates and cultured in complete DMEM medium until reaching ~70–90% confluency. rKSHV.219 viral supernatant was thawed rapidly in a 37°C water bath and immediately placed on ice. For infection, 1 mL of viral supernatant with 8 µg/mL polybrene was added per well and the plates were centrifuged at 1,200 × g for 90 minutes at room temperature using a swinging-bucket rotor. After centrifugation, plates were returned to a 37°C incubator for 6–16 hours. The medium was then replaced with 2 mL of fresh complete DMEM. Forty-eight hours post-infection, the infected cells were selected and maintained with 1 µg/mL puromycin.

### Luciferase assays

HCT116 p53 KO, BC3 and BJAB219 cells were co-transfected in 24-well plates with a total of 500 ng DNA per well using PolyJet transfection reagent (SL100688; SignaGen Laboratories). Each well received 150 ng of PURPL wild-type (WT) or PURPL mutant firefly luciferase reporter plasmid, 5 or 15 ng of the indicated expression plasmid unless stated otherwise, and 50 ng of the internal control plasmid pGL4.74[hRluc/TK] (Renilla luciferase; #E6921, Promega). For selected experiments, the empty luciferase vector pGL4.11[luc2P] (#<u>E6661</u>, Promega) was used as a control plasmid. Total DNA amounts were kept constant at 500 ng per well by supplementing with pBluescript SK(-) empty vector as needed. At 48 hours post-transfection, cells were harvested and luciferase activity was measured using a TECAN Infinite M1000 Multilabel Microplate Reader. Relative luciferase units (RLU) were calculated by dividing firefly luciferase activity of the tested reporter plasmid by Renilla luciferase activity of the internal control.

### Transductions

The shRNA vectors were based on the GIPZ lentiviral backbone. The non-silencing control shRNA vector was a kind gift from Yosef Shaul and the shRNAs targeting p73 were purchased from Horizon Discovery as a set of five plasmids, clone IDs; V3LHS_330454 named as sh-p73 (1), V2LHS_69894 named as sh-p73 (2), V3LHS_330455 named as sh-p73 (3), V3LHS_330453 named as sh-p73 (4) and V2LHS_181826 named as sh-p73 (5). The shRNA lentiviruses were prepared in HEK293T cells. The cells were transfected with 2 μg of respective shRNA plasmid, 2 μg psPAX2 (Plasmid #12260, Addgene) and 1 μg pMD2.G (Plasmid #12259, Addgene) packaging plasmids. Forty-eight hours after transfection, the supernatant containing lentivirus was collected and centrifuged at 1200 rpm for 5 minutes at 4°C and filtered through a 0.45 μm filter. The filtered lentiviral supernatant was then added to BJAB cells in the presence of 4 μg/mL polybrene and incubated. After 4 hours, the media was replaced with fresh media. Forty-eight hours after transduction, BJAB cells were selected using 1 μg/mL puromycin, as the GIPZ backbone encodes puromycin resistance. BJAB cells were also transduced with pLIX403 or pLIX403_LANA lentiviral vectors using the same protocol.

### RNA isolation and RT-qPCR

Total RNA was isolated using RNeasy Kit for RNA Purification (Qiagen), the lysate was homogenized using Qiashredder kit (Qiagen) and reverse transcription was performed using a high-capacity cDNA reverse transcription kit (Applied Biosystems). The qPCR was performed using Fast SYBR green master mix (Applied Biosystems) and analyzed with a CFX96 Touch real-time PCR detection system (Bio-Rad). GAPDH was used as an internal control (reference gene) for mRNA expression. All primers used are listed in S1 Table.

### Flow cytometry

FACS was performed by annexin V-FITC apoptosis detection with PI staining kit (BioLegend) and then assessed using flow cytometry quantification as per manufacturer’s protocol. Flow cytometry analysis was performed on Gallios flow cytometer (Beckman Coulter) and all data were analyzed by FlowJo software (BD Biosciences).

### RNA-sequencing analysis

BAM files from Journo et al 2018 ^43^ were uploaded onto the UCSC human genome browser ^61^.

### ChIP-sequencing analysis

Available ChIP-seq data for p53 and p73 (GEO: GSE15780)^47^ and for p63 (GSE135257)^48^ were reanalyzed using Partek™ Genomics Suite™ software to determine the association of the proteins with PURPL promoter. FASTQ files were aligned to the human genome (hg19) using Bowtie2, and BAM files were visualized using the chromosome view option.

## Acknowledgments

We would like to thank David Root, Yosef Shaul and Moshe Oren for kindly providing the plasmids, Bert Vogelstein, Yosef Shaul, Jae U Jung, Ronit Sarid, Jeffrey Vieira, Richard F. Ambinder, and S. Diane Hayward for cell lines, and Yosef Shaul for critically reading the manuscript.

## Author Contributions

Conceived and designed the experiments: S.S., G.J., D.M., V.G., M.S. Performed the experiments: S.S., G.J., D.M, Analyzed the data: S.S., G.J., D.M., V.G., M.S. Generated tools & reagents: S.S., G.J., D.M., V.G, Wrote the paper: S.S., M.S.

