## Supplementary figures for "KSHV upregulates the long non-coding RNA PURPL via p73 to suppress p53"

**A**

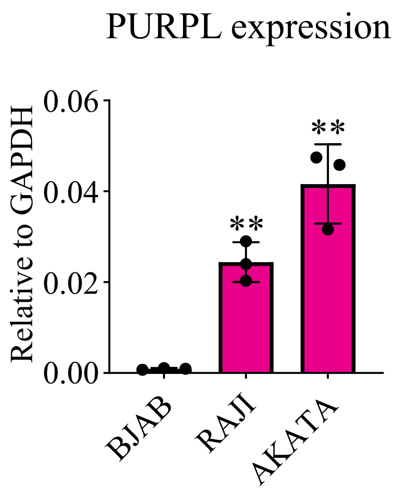

**Figure S1. PURPL expression in EBV-positive B-cell lines.** (A) RT-qPCR analysis of PURPL RNA levels in BJAB (EBV-negative), RAJI, and AKATA (EBV -positive) cells. Gene expression was normalized to GAPDH. Experiments were performed in three biological replicates (n = 3), and data are represented as means ± SD. One-tailed T-tests were performed.

### Figure S1

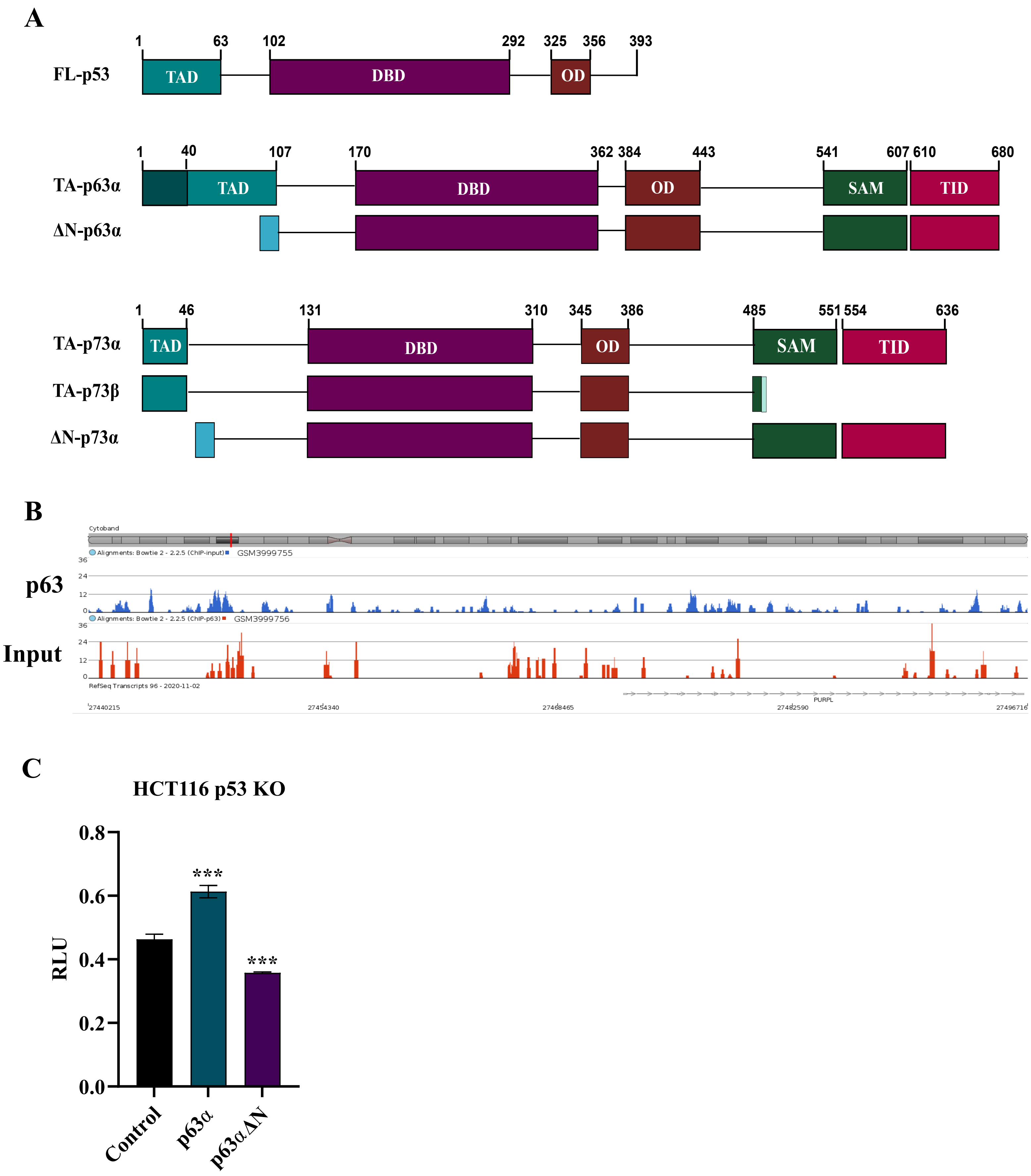

**Figure S2. p53 family domain structure and regulation of the PURPL promoter by p63.** (A) Schematic representation of p63α, ΔNp63α, p73α, p73β and ΔNp73α conserved domains. (B) Re-analysis of publicly available p63 ChIP-seq data (GEO: GSE135257) generated in ME-180 cells was performed using Partek™ Genomics Suite ™, and binding profiles were visualized using the chromosome view option. (C) HCT116 p53 KO cells were transfected with luc-PURPL promoter together with expression vectors for p63α or ΔNp63α (15 ng each), and dual luciferase reporter assay was performed. Experiments were performed in three biological replicates (n = 3) and data are represented as means ± SD. Luciferase data are presented as relative luciferase units (RLU) normalized to Renilla luciferase internal control.

**A**

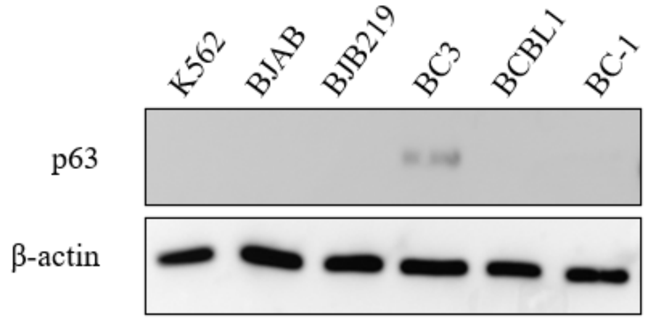

**Figure S3. p63 protein expression in uninfected and KSHV-infected B-cell lines.** Western blot analysis of p63 protein levels in K562 and BJAB (virus -negative), BJAB.219 (de novo KSHV-infected), BC3 and BCBL1 (chronically KSHV -infected), and BC -2 (chronically infected with KSHV and EBV) cells. β-actin was used as a loading control.

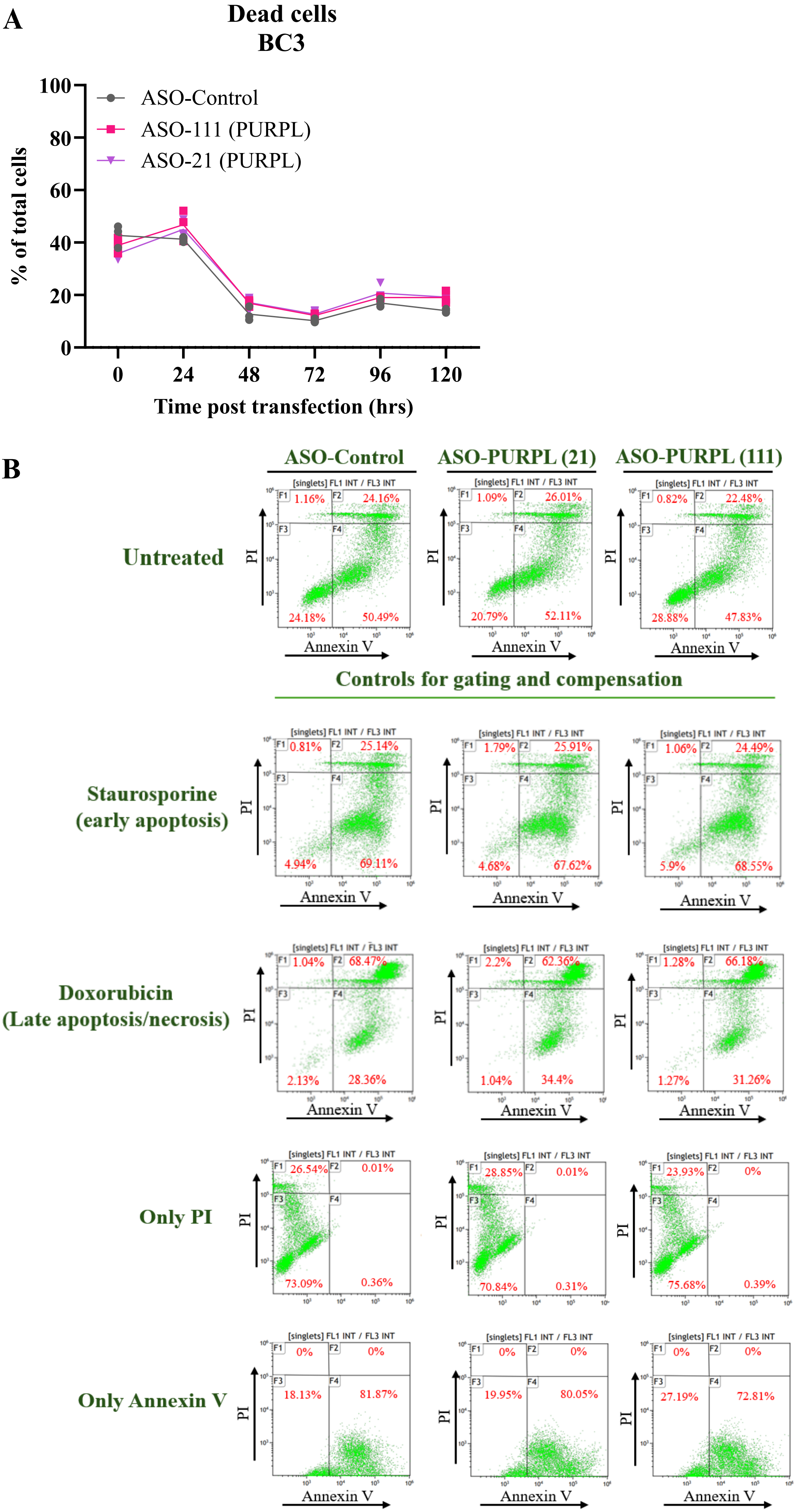

A
Dead cells
BC3
% of total cells
ASO-Control
ASO-111 (PURPL)
ASO-21 (PURPL)
Time post transfection (hrs)
B
ASO-Control
ASO-PURPL (21)
ASO-PURPL (111)
Untreated
1.16% 24.16% 24.18% 50.49%
1.09% 26.01% 20.79% 52.11%
0.82% 22.48% 28.88% 47.83%
Controls for gating and compensation
Staurosporine
(early apoptosis)
0.81% 25.14% 4.94% 69.11%
1.79% 25.91% 4.68% 67.62%
1.06% 24.49% 5.9% 68.55%
Doxorubicin
(Late apoptosis/necrosis)
1.04% 68.47% 2.13% 28.36%
2.2% 62.36% 1.04% 34.4%
1.28% 66.18% 1.27% 31.26%
Only PI
26.54% 0.01% 73.09% 0.36%
28.85% 0.01% 70.84% 0.31%
23.93% 0% 75.68% 0.39%
Only Annexin V
0% 0% 18.13% 81.87%
0% 0% 19.95% 80.05%
0% 0% 27.19% 72.81%
PI
Annexin V
[singlets] FL1 INT / FL3 INT

**Figure S4. PURPL knockdown does not alter apoptosis in BC3 cells.** (A) BC3 cells transfected with ASO-Control or ASO-PURPL (ASO-21 and ASO-111) were counted every 24 h via an automated cell counter to evaluate the percentage of dead cells. The cell count assay was performed in three biological replicates ($n = 3$), and data are represented as means ± SD. (B) Flow cytometry analysis of apoptosis in BC3 cells transfected with the same ASOs shown in panel A. Annexin V/PI staining plots are shown. Single-staining controls (Annexin V only and PI only) were used to establish compensation and gating parameters. Staurosporine treatment served as a positive control for early apoptosis, and doxorubicin treatment served as a positive control for late apoptosis/necrosis. Untreated cells are shown as baseline controls.

**Supplementary Table 1 - Primer sequences**

| | |
|---|---|
| **Antisense oligos** | |
| ASO-NC5 (named as ASO-Control) | 52MOErG/i2MOErC/ i2MOErG/ i2MOErA/ i2MOErC/TATACGCGCA/ i2MOErA/ i2MOErT/ i2MOErA/ i2MOErT/32MOErG |
| ASO-21 | 52MOErG/ i2MOErG/ i2MOErT/ i2MOErT/ i2MOErC/CTGCTACATA/ i2MOErG/ i2MOErC/ i2MOErA/ i2MOErC/32MOErC |
| ASO-111 | 52MOErT/ i2MOErA/ i2MOErG/ i2MOErG/ i2MOErC/ACACGGGCAC/ i2MOErA/ i2MOErT/ i2MOErT/ i2MOErG/32MOErA |
| **Primer list for RT-qPCR** | |
| GAPDH- F | 5’-CATCATCTCTGCCCCCTCT -3’ |
| GAPDH- R | 5’-GGTGCTAAGCAGTTGGTGGT-3’ |
| PURPL isoform 5- F | 5'-TGATTGGAAATGTGTGCTTCCC -3' |
| PURPL isoform 5- R | 5'-AGAACCCTGTATGTGGACTGTG -3' |
| P73- F | 5’-GCGTGGAAGGCAATAATCTCT -3’ |
| P73- R | 5’-GTTGTACAGGATGGTGGTGAA -3’ |
| LANA- F | 5’- GAAGTTGTAGGAAACGAAACAGGT-3’ |
| LANA- R | 5’-ACACTGTGGGACTTCCAGGTATAG -3’ |
| **Primers used for cloning** | |
| PURPL promoter -F | 5’-AAGGAAACCTCAAAACGTGAAGGAC-3’ |
| PURPL promoter -R | 5’-AAAGCAAGAGGGGTTCCTGCTAC-3’ |
| Q5SDM -F | 5’- TAGATC TGCCCACAGTGGACTGGAG -3’ |

**Supplementary Table 2- Nucleotide substitutions introduced into the p53 binding site of the PURPL promoter**

| **Mutation of the p53 binding site in the PURPL promoter** | |
|---|---|
| Wild-Type | TGAGGGAACCTGCCCAGGGCTTGTCTGGGCATGCCCACAGTGGACTG GAGCCTGACATGCT CACTGGGGCAA |
| Mutant | TGAGGGAACCTGCCCAGGGGACGTCTAGATCTGCCCACAGTGGACTG GAGCCTGACATGCT CACTGGGGCAA |

Substituted nucleotides are indicated in red.
